# Functional diversity in a tritrophic system: Effects on biomass production, variability, and resilience of ecosystem functions

**DOI:** 10.1101/431981

**Authors:** Ruben Ceulemans, Ursula Gaedke, Toni Klauschies, Christian Guill

## Abstract

Diverse communities can adjust their trait composition to altered environmental conditions, which may strongly influence their dynamics. Previous studies of trait-based models mainly considered only one or two trophic levels, whereas most natural system are at least tritrophic. Therefore, we investigated how the addition of trait variation to each trophic level influences population and community dynamics in a tritrophic model. Examining the phase relationships between species of adjacent trophic levels informs about the degree of top-down or bottom-up control in non-steady-state situations. Phase relationships within a trophic level highlight compensatory dynamical patterns between functionally different species, which are responsible for dampening the community temporal variability. Furthermore, even without trait variation, our tritrophic model always exhibits regions with two alternative states with either weak or strong nutrient exploitation, and correspondingly low or high biomass production at the top level. However, adding trait variation increased the basin of attraction of the high-production state, and decreased the likelihood of a critical transition from the high-to the low-production state with no apparent early warning signals. Hence, our study shows that trait variation enhances resource use efficiency, production, variability, and resilience of entire food webs.

## 1 Introduction

Functional diversity has proven to be important for linking community structure to ecosystem functions such as biomass production and resource use efficiency (Hooper et al., 2005; Tilman et al., 2006; Worm et al., 2006; Schneider et al., 2016). Our understanding of the multifaceted impact of functional diversity on ecosystem functioning, and on the dynamics of populations and communities, has been greatly advanced by adopting a trait-based point of view (Hillebrand and Matthiessen, 2009; Krause et al., 2014). In particular, functional traits link morphological, physiological or phenological features of a species to a certain community or ecosystem function (Violle et al., 2007). A prevalent example is simply body size, which is related to several functions such as growth (larger organisms tend to grow slower), prey preference (predators tend to be larger than their prey), or nutrient uptake (larger cells have higher nutrient demands) (Weithoff, 2003; Brown et al., 2004; Brose et al., 2006; Litchman et al., 2007). Trait-based models of simple food web modules have facilitated detailed mechanistic understanding of dynamics observed in the laboratory (Ellner and Becks, 2011) and in the field (Tirok and Gaedke, 2010). For example, observed anti-phase predator-prey cycles between zooplankton and algae have been attributed to the co-occurrence of fast-growing undefended and slow growing, well defended prey phenotypes (Yoshida et al., 2003; Becks et al., 2010).

However, such trait-based models have mainly been restricted to describing trait variation on one or two trophic levels (Abrams and Matsuda, 1996; Litchman and Klausmeier, 2008; Tirok and Gaedke, 2010; Erbach et al., 2013). Likewise, only up until recently, empirical studies on functional diversity have been limited to considering trait variation in only autotrophs (plants or algae) (Duffy, 2002; Gamfeldt et al., 2015), or both autotrophs and herbivores (Steiner et al., 2005; Filip et al., 2014), with few exceptions (Rasher et al., 2013). This strongly contrasts with the fact that natural food webs are in general complex multitrophic networks (Digel et al., 2014). Focusing only on direct, bitrophic predator-prey interactions neglects the intricate effects of more complex, partly indirect interactions spanning multiple trophic levels, such as trophic cascades (Levine et al., 2017). These multi-trophic effects may be very important factors affecting the relevant ecosystem functions (Peet et al., 2005; Nakazawa, 2011; Golubski et al., 2016). For instance, the total number of trophic levels may strongly influence the efficiency of nutrient exploitation (Wang and Brose, 2018). In addition, as predation is an important factor in many food webs, trait variation on the predator level is expected to have an important influence on ecosystem functioning (Tilman et al., 2014; Gamfeldt et al., 2015; Schneider et al., 2016). Hence, including additional trophic levels with functional diversity is a very natural step towards improving the accuracy and descriptive power of trait-based models.

We developed a tritrophic model to study the effects of trait variation at all trophic levels on food web dynamics. Particularly, the dynamics of a simple tritrophic linear food chain will be compared to a tritrophic food web where prey species are either defended or undefended, and predator species are either generalist or specialist feeders. Trade-offs between these traits are explicitly built in such that defended prey have a lower growth rate, and specialist feeders have a lower half-saturation constant to allow for efficient feeding at low prey densities (Tirok and Gaedke, 2010; Coutinho et al., 2016; Van Velzen and Gaedke, 2017). Our model structure allows for a gradual increase of the trait differences between the species at each trophic level, from a simple linear chain up to a fully separated food web with maximal trait differences (Fig. 1). As the trait differences increase, the species will fulfill increasingly different functions; in this way, we are able to link trait differences to functional diversity.

**Figure 1:**
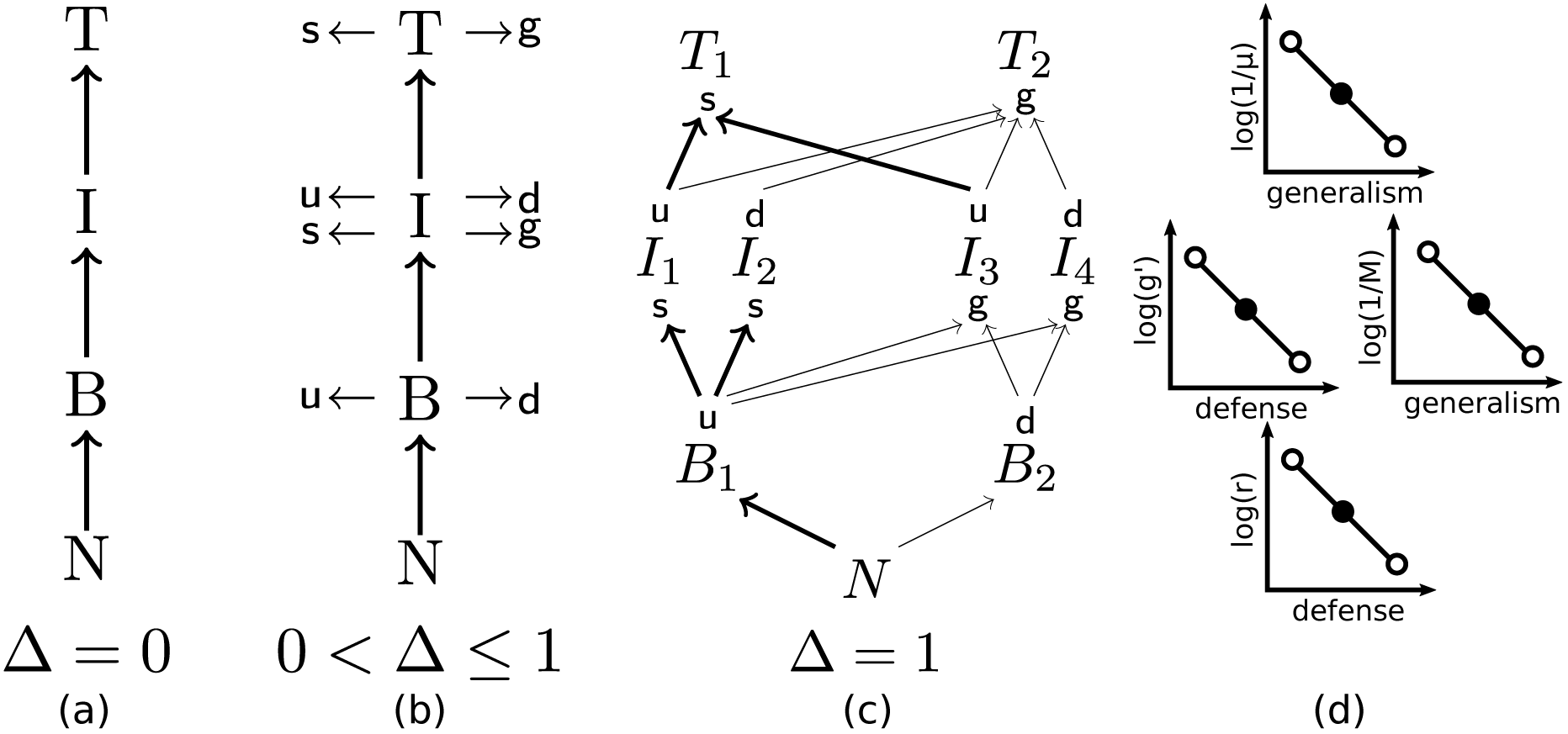
(a) A simple linear tritrophic chain, where nutrients N are taken up by a basal species B, which is grazed by an intermediate species I, which in turn is consumed by a top species T. (b) Gradually introducing trait variation, where species can be undefended (u) or defended (d) against predators, and/or specialist (s) or generalist (g) consumers, starting from a linear chain. (c) Maximally trait-separated food web model. The basal and top species have only one trait, but the intermediate species have two. The thickness of the arrows indicates the intensity of the trophic interaction, reflecting that specialist consumers can exploit their limited resources spectrum more efficiently. (d) Schematic shape of the trade-off curves for the top species (top row), intermediate species (middle row), and basal species (botton row). The solid circles indicate that for ∆ = 0, all species on a given trophic level have the same trait values, whereas the open circles demonstrate how the trait values between the species differ as ∆ is increased to one.

We use the tritrophic model to investigate how an increasing degree of trait variation affects:

- the production of the system,
- the efficiency of the energy transfer towards the higher trophic levels,
- the temporal biomass variability at the population and community level, and
- the dynamic properties and the resilience of alternative stable system states.

Our results provide theoretical evidence that trait variation has a significant impact on all of these properties. To our best knowledge, we present the first systematic, multi-trophic study which mechanistically explains such patterns and explicitly discusses their relevance to ecosystem functions and stability.

## 2 Methods

We developed a tritrophic model where basal species are consumed by intermediate species, which in turn are consumed by top species. The species biomass densities are denoted by respectively *B*, *I*, and *T*. In addition, the uptake of a limiting nutrient with concentration *N* (in this case nitrogen) by the basal species is modeled explicitly. We assume a chemostat environment, which causes all nutrients and biomass of species to be washed out at an equal rate, *δ*, the dilution rate. The washed out volume is replaced by new medium rich in nutrients.

### 2.1 Model equations

As in (Tirok and Gaedke, 2010; Bauer et al., 2014), we include two relevant functional traits. The prey species *B* and *I* may be defended against predation: specifically, there will be defended (d) and undefended (u) species. Investing in a defense strategy requires sacrificing a certain amount of resources which could have otherwise been put into growth. Hence, the defended species have a lower growth rate than the undefended species, but are rewarded by being insusceptible to certain consumers. The consumer species *I* and *T* are able to specialize feeding on a certain prey species, leading to specialist (s) and generalist (g) species. Here, the generalist consumer species are able to consume all species on the trophic level below. In contrast, the specialist species are only able to consume the undefended prey species, but they are able to exploit low food densities at a higher rate, reflected in a lower half-saturation constant.

Representing each possible trait combination on all trophic levels by one species leads to a food web with two basal species, four intermediate species and two top species (Fig. 1c). In order to write down the equations compactly, the following equivalence is explicitly stated:

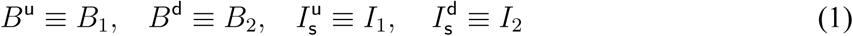

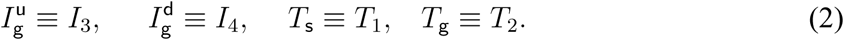

In their most general form, the equations used have the following shape:

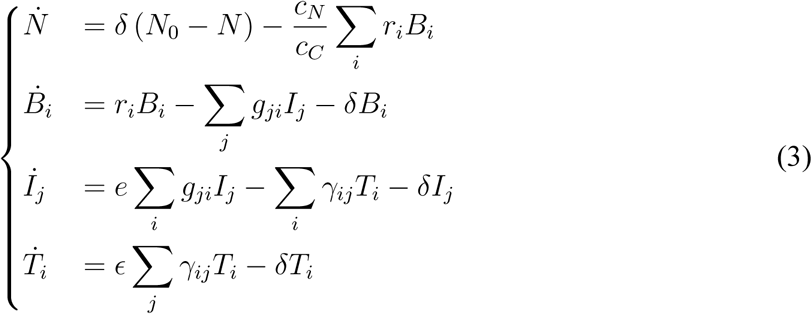

with *i ∈* {1, 2}, *j ∈* {1, 2, 3, 4}, where *N*_0_ denotes the incoming nutrient concentration. Following typical experimental conditions, we assume nitrogen as the limiting nutrient (*N*). Hence, the nutrients are measured in nitrogen concentration, as compared to carbon for biomass, therefore, the nitrogen-to-carbon weight ratio (*c_N_* /*c_C_*) is required to scale the basal (*B_i_*) growth terms. Moreover, the basal growth rate *r_i_* is described by a Monod function (Monod, 1950; Tilman et al., 1982), with maximum growth rate 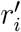 and a nutrient-uptake half-saturation constant *h_N_*. The intermediate and top species have a generalized Holling-type-III functional response, with maximum growth rates 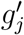 and 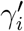, half-saturation constants *M* and *μ*, and Hill coefficients *h* and *η*, respectively (Williams and Martinez, 2004; Kalinkat et al., 2013). This means:

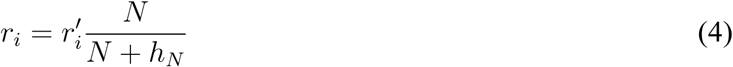

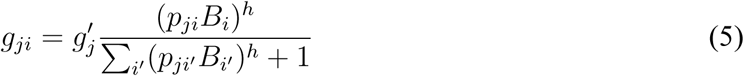

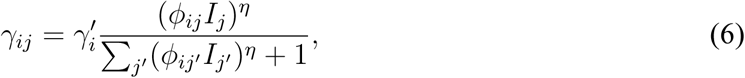

and,

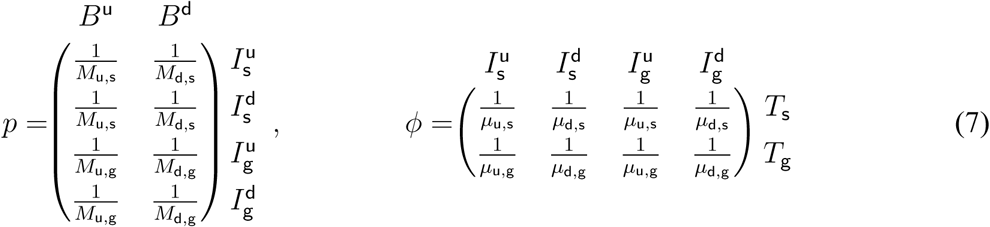

such that e.g. *M*_u,s_ indicates the half saturation constant of the undefended species being grazed by the specialist species, etc.

Finally, our model includes a parameter, ∆, which explicitly controls the species’ trait values. Abstract traits such as defense and specialization are linked to concrete and measurable parameters describing the species’ interactions. For the basal species, their maximal growth rate 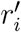 is linked to their position on the defense axis (Fig. 1d). The intermediate species have two trait values: defense is again linked to their maximal growth rate 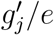, and the half saturation constant *M_i_* is determined by their degree of specialization; both of these traits affect the overall growth rate of the intermediate consumers. The top species have only one trait, specialization, which is linked to their half saturation constant *μ_i_*. As will be shown below, the equations are parametrized in a way such that for ∆ = 0 the linear chain system, where all species per trophic level are functionally identical, will be described (Fig. 1a). As ∆ is increased, the system changes in a continuous way, where some prey species gradually become more and more defended (Fig. 1b), such that they can be preyed on less and less by the specialist species. In addition, the specialist species are gradually able to feed more efficiently on the undefended species. Finally, for ∆ = 1 the trait differences are maximal, as is the case in Fig. 1c: the specialist species are not able to feed on the defended prey anymore.

The parameter values are set to vary logarithmically with ∆. This implies that parameter changes occur proportional to the starting value in both directions, since 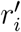 and 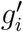 appear as linear factors in the differential equations. For consistency, the elements of *p* and *ϕ* are also varied logarithmically. Concretely, this means that:

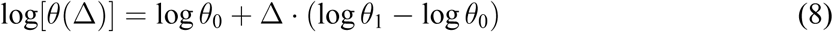

where θ is 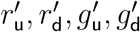 or any or *p_ij_* or *ϕ_ji_*. In this way, Δ = 0 implies *θ* = *θ*_0_ such that all the trait values are equal in the following manner:

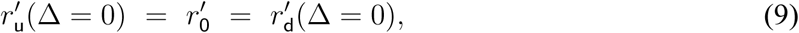

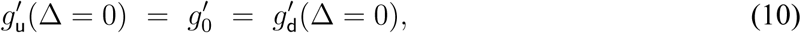

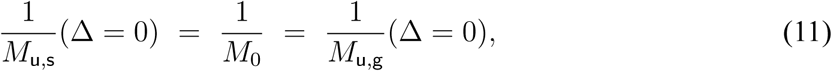

and similarly for the other elements of *p* or *ϕ*. We define the parameter values *θ*_0_ of the ∆ = 0 system as arithmetic averages of the extreme values *θ*_1_ in the ∆ = 1 system, on a logarithmic scale, e.g.:

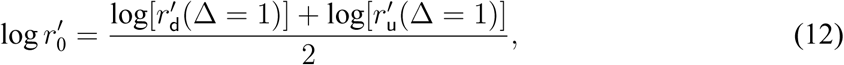

and similarly for the other parameters. These extreme values are shown in Table 1.

**Table 1:** Standard parameter values used in this study when the trait differences are maximal, i.e., ∆ = 1.

|  |  |
| --- | --- |
| Body mass ratio between adjacent trophic levels | $m_I/m_B = m_T/m_I = 10^3$ |
| Allometric scaling exponent | $\lambda = -0.15$ |
| Inflow nutrient concentration | $N_0 = 1120 \mu\text{g N/l}$ |
| Dilution rate | $\delta = 0.055$ |
| Nutrient half-saturation const. of $B$ | $h_n = 10 \mu\text{g N/l}$ |
| Nitrogen to carbon ratio of $B$ | $c_N/c_C \approx 0.175$ |
| $B^u$ max. growth rate | $r'_1 = 1$ |
| $B^d$ max. growth rate | $r'_2 = 0.66$ |
| $I$ conversion efficiency | $e = 0.33$ |
| $I^u$ max. grazing rate | $g'_{1,3} \approx 1.06$ |
| $I^d$ max. grazing rate | $g'_{2,4} \approx 0.70$ |
| $I_s$ half-saturation const. | $M_{1,2} = 300 \mu\text{g C/l}$ |
| $I_g$ half-saturation const. | $M_{3,4} = 600 \mu\text{g C/l}$ |
| $T$ conversion efficiency | $\epsilon = 0.33$ |
| $T$ max. grazing rate | $\gamma'_{1,2} \approx 0.38$ |
| $T_s$ half-saturation const. | $\mu_1 = M_{1,2} = 300 \mu\text{g C/l}$ |
| $T_g$ half-saturation const. | $\mu_2 = M_{3,4} = 600 \mu\text{g C/l}$ |

As the logarithm of 0 is undefined, this requires the elements of *p* and *ϕ* related to the defended-specialist species’ interactions for ∆ = 1 to be nonzero; in this case 10^*−*4^ was taken. Note also that the set of 9 equations in Equation (3), when the species on each trophic level are exactly equal (∆ = 0), is mathematically equivalent to a linear chain system with 4 equations, up to a slight parameter transformation. Specifically, the 9-equation food web system corresponds to a 4-equation food chain by setting *M →* 2^(*h−*1)/*h*^M and *μ →* 4^(*h−*1)/*h*^μ. For details of the derivation, see Appendix A.

### 2.2 Model parametrization and analysis

In order to decrease the number of free parameters, and simultaneously increase the realism of the model, the species’ growth rates were scaled allometrically to their body mass (Brose et al., 2006; De Castro and Gaedke, 2008):

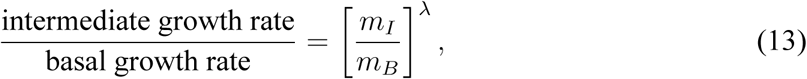

with body masses *m* and the exponent *λ* given typical for planktonic systems (Moloney and Field, 1989) (Table 1). The same relationship holds true for the ratio between the maximum growth rates of the intermediate and the top species.

This model was developed as a chemostat model, with an eye towards potential experimental application. Chemostat experiments have been very successful in identifying and understanding ecological and evolutionary interactions of planktonic (Fussmann et al., 2005), and many other microbiological systems (Elena and Lenski, 2003). In such experiments, many factors influencing dynamics in question, such as nutrient supply, light supply, temperature, etc., are kept constant and/or closely monitored. This procedure greatly facilitates observation of the interactions of interest between species in the chemostat. For this reason, extra care was taken to have empirically motivated and realistic values of the remaining model parameter values (Table 1). Specifically, the parameter values we use are representative for planktonic chemostats. However, this does not mean that our results apply only to planktonic systems. In fact, as we show in Appendix B, similar results are obtained when the model parametrization is more adapted towards terrestrial food webs, for example.

To get a better understanding of how much certain values of ∆ sets the species on the three trophic levels apart, we here consider a few exemplary cases. At ∆ = 0, the varied parameters are identical (and so are the species), while at ∆ = 0.2, maximal growth and grazing rates of the undefended species are 9% higher than those of defended species, and half saturation constants of generalists are 15% higher than generalists. At ∆ = 0.5, the differences are 23% and 42%, respectively; at ∆ = 1 they are 50% and 100%.

For simplicity and to reduce the dimensionality of the system somewhat, in the rest of the text it will be assumed that

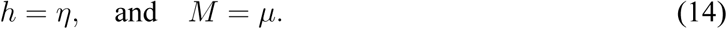

Hence, *h* will denote the Hill exponent, and *M* the half-saturation constant, of the functional response between both the first and the second, and between the second and the third trophic level (implying *p* = *ϕ*). Additional narrowing of parameter ranges was achieved by requiring coexistence of the species in both the chain and the maximally separated food web.

To characterize the differences between the different attractors, the different phase relationships between predator-prey pairs were investigated. These phase relationships were obtained by calculating the Discrete Fourier Transform (DFT) of the simulated time series. Due to the non-sinusoid shape of the biomass oscillations, a signal with only a single frequency will generate an infinite amount of peaks in the frequency spectrum. These are necessarily multiples of the original frequency *f*, and the height of the peaks will scale as 1/*f* (Bracewell, 1999). This means they are easily identified in the frequency spectra when shown on a log-scale, by the linear decay in peak height.

The solutions of the differential equations presented were obtained numerically in C using the SUNDIALS CVODE solver (Hindmarsh et al., 2005), with relative and absolute tolerances of 10^*−*10^. Output data were studied using Python and several Python packages; in particular NumPy, SciPy and Matplotlib (Van Der Walt et al., 2011; Hunter, 2007).

## 3 Results

Firstly, we compare the biomass dynamics of the linear chain to the dynamics of the maximally trait-separated food web, where trait differences within each trophic level are maximal. Secondly, we study certain properties of the system, such as the temporal variability of population and community biomasses, and the relative abundances of species, while gradually increasing the amount of standing trait variation at each trophic level from a linear food chain to the maximally trait-separated food web.

The amount of trait variation is described by ∆ ranging continuously from 0 to 1. When ∆ = 0 we describe the linear chain without trait variation, and when ∆ = 1 we describe the maximally trait-separated food web. This fully separated food web consists of defended and undefended prey species, which are being preyed upon by generalist and/or specialist predator species (Fig. 1c). The benefits and costs of the different offense-defense strategies are linked to each other through predefined trade-offs (Methods). The defended species have a lower growth rate than the undefended species, but in turn, they are not preyed upon by the specialist species of the next trophic level in the fully separated web. Similarly, the specialists, while unable to prey on the defended species, are able to graze the undefended species more efficiently at low prey concentrations than their generalist counterparts.

Our results are first summarized schematically in Fig. 2, subsequent mechanistic details are presented in the sections and figures below. We observe two alternative stable states with low vs. high total production (State 1 and 2 in Fig. 2) in both the linear chain (low trait variation) and the maximally separated food web (high trait variation). In the low-production state, the high mean concentration of free nutrients corresponds to a low amount of total biomass and consequently, a low total production. In the high-production state, in contrast, the low mean nutrient concentration implies that most of the nutrients are stored in the biomass which implies a high total production. Note that at equilibrium, the total amount of nutrients in the system is always constant because the chemostat model’s dilution rate *δ* is constant for all species.

**Figure 2:**
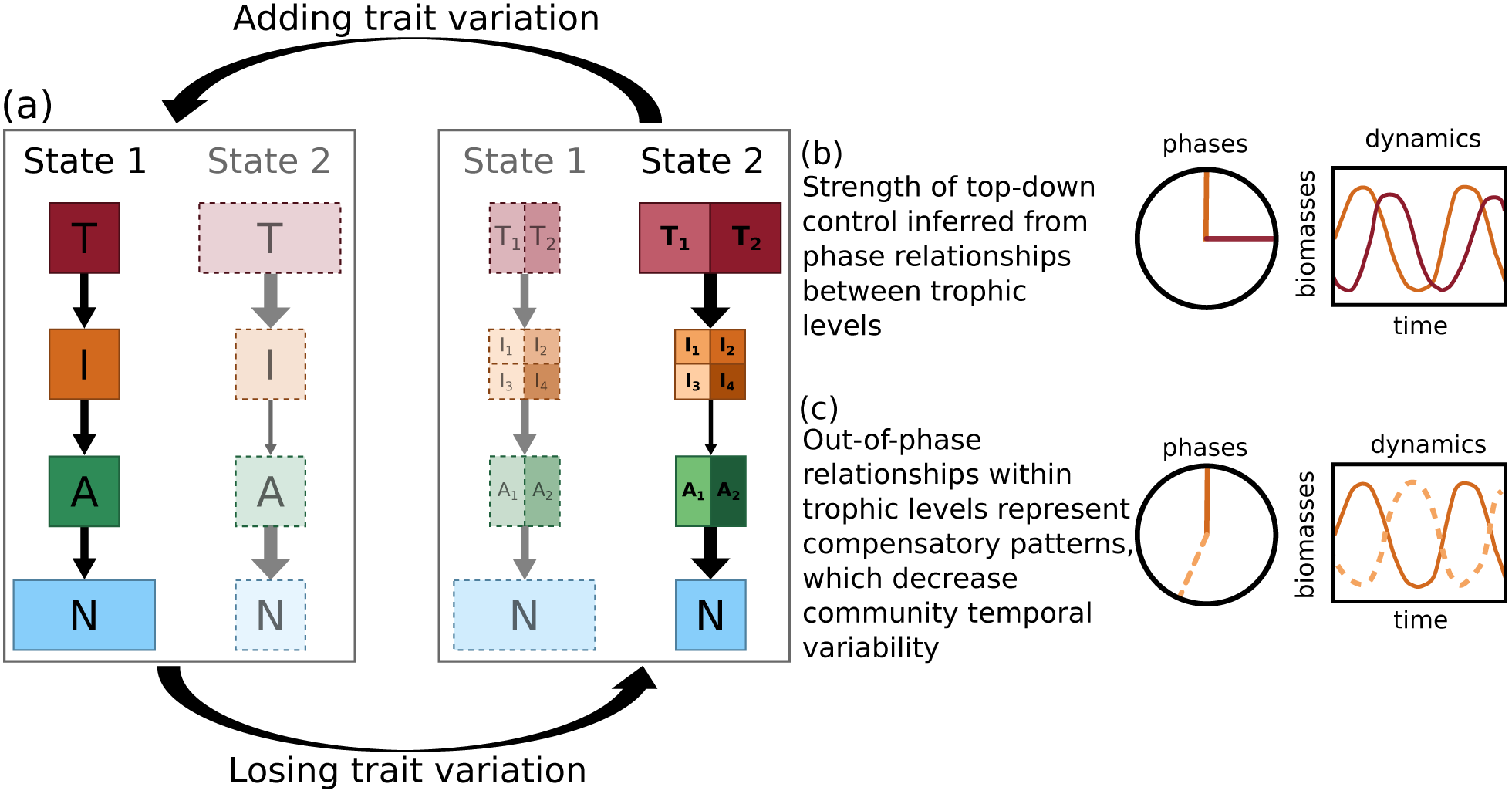
Schematic overview of our results. (a) The model system (with A autotrophs, I intermediate consumers and T top predators) always exhibits two alternative stable states, State 1 (low-production) and 2 (high-production), for both low and high amounts of trait variation. As trait variation is added, the system tends toward the high-production state with the top-heavy biomass pyramid (solidly drawn states have a larger basin of attraction than grayed out states). (b) The phase differences between predators and their prey inform about the intensity of top-down control in the system as indicated by arrow width in (a), i.e. ¼-lag cycles indicate strong coupling between predator and prey. (c) Within trophic level out-of-phase cycles indicate compensatory dynamical patterns, where the different species exploit different temporal niches, and hence, reduce the community temporal variability.

In the food chain without trait variation (left part of the biomass pyramids in Fig. 2), the population-level biomass dynamics for the low-production state (Fig. 3a) exhibit pronounced predator-prey cycles, while the high-production state exhibits slower cycles with lower amplitudes (Fig. 3b). The respective phase relationships of these oscillations (right part in Fig. 2, Fig. 3c & 3d) inform about the ecological mechanism behind the two different states (for details, see section 3.1). In the low-production state, fast cycles with high amplitudes occur due to the strong coupling between adjacent trophic levels. Such a strong interaction between predators and their prey is indicated by the quarter-cycle phase lags (henceforth referred to as ¼-lag cycles) (Fig. 3c). In the high-production state the top and intermediate level still exhibit ¼-lag cycles, but the phase difference between the intermediate and basal level is significantly larger (Fig. 3d). This offset in the phase-relationship indicates that the top-down control over the intermediate level is so strong in the high-production state that the intermediate level’s dynamics are less closely coupled to the basal level than in the low-production state. The basal level is then free to fully exploit the available nutrients.

**Figure 3:**
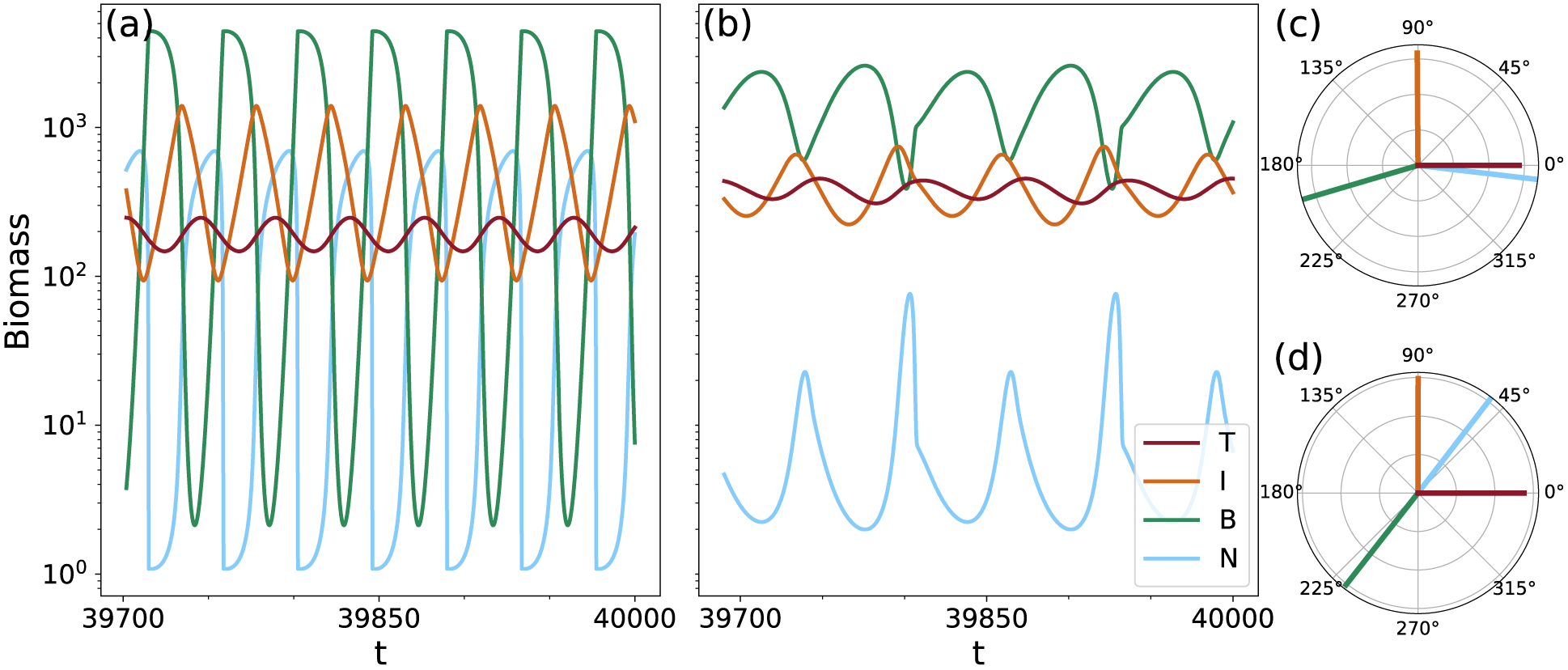
Biomass and nutrient dynamics on the two different states for the tritrophic chain for h = 1.1 (= η), and their corresponding phase relationships. (N = nutrients, B = basal species, I = intermediate species and T = top species). The relative phases of the low-production state shown in (a) (mean nutrient level ≈ 250 μg N/l) are plotted in panel (c). The phases of the high-production state shown in (b) (mean nutrient level ≈ 10 μg N/l) are plotted in panel (d). In both cases the phases relative to the top species are shown.

The decoupling of the intermediate and basal level results in a lower temporal variability, especially at the basal level, and hence, reduces times of strong basal suppression during which the nutrients can almost reach their capacity as observed in the low-production state. In the high-production state, the overall higher primary production combined with the lower temporal variability between the basal and the intermediate level enhances the energy transfer through the food chain and results in a top-heavy biomass pyramid.

When the food chain becomes a food web by adding trait variation (right part of the biomass pyramids in Fig. 2), the biomass dynamics of the low-production (Fig. 4a) and high-production state (Fig. 4b) as well as their respective phase-relationships (Fig. 4c - 4f) become more complex because a slow and a fast timescale underlie the oscillations (see Section 3.1 for details). Information regarding intensity of top-down control is only deduced from the phase-relationships of the oscillatory mode that explains most of the observed variation, i.e. the fast timescale of the low-production state (Fig. 4d) and the slow timescale of the high-production state (Fig. 4e). Similar to the food chain without trait variation, the strong top-down control by the top level and subsequently, the decoupling of the intermediate and basal level in the high production state again results in lower temporal variability, a temporally more balanced nutrient use, and a more efficient energy transfer towards the top level. Importantly, the high-production state becomes more likely than the low-production state with increasing trait variation, i.e., its basin of attraction increases, making it more resilient against external disturbances (for details, see Fig. 5 and Section 3.2).

**Figure 4:**
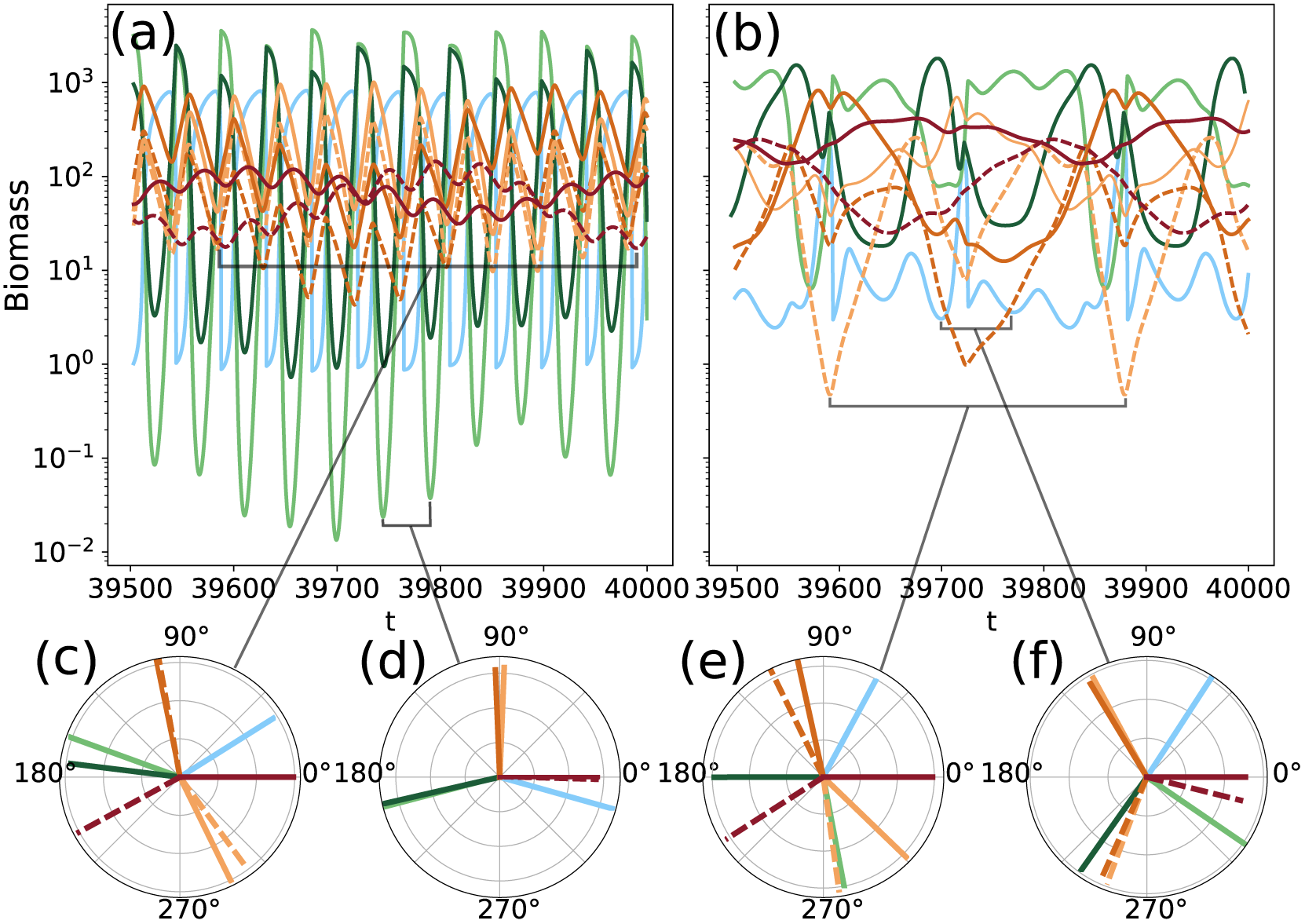
The dynamics of the maximally separated food web (see Figure 1c for structure and species names), for h = 1.05 (= η). Figure (a) and (b) show the biomass time series on the low-and high-production state, respectively. The phase relationships (relative to T_g_) of the two main temporal modes on the states are shown in panels (c), (e) (slow) and (d), (f) (fast). (See Figure 5 and its explanation in the text for why the chosen value of h = 1.05 is different from the one used to compare the two states on the chain in Figure 3.)

**Figure 5:**
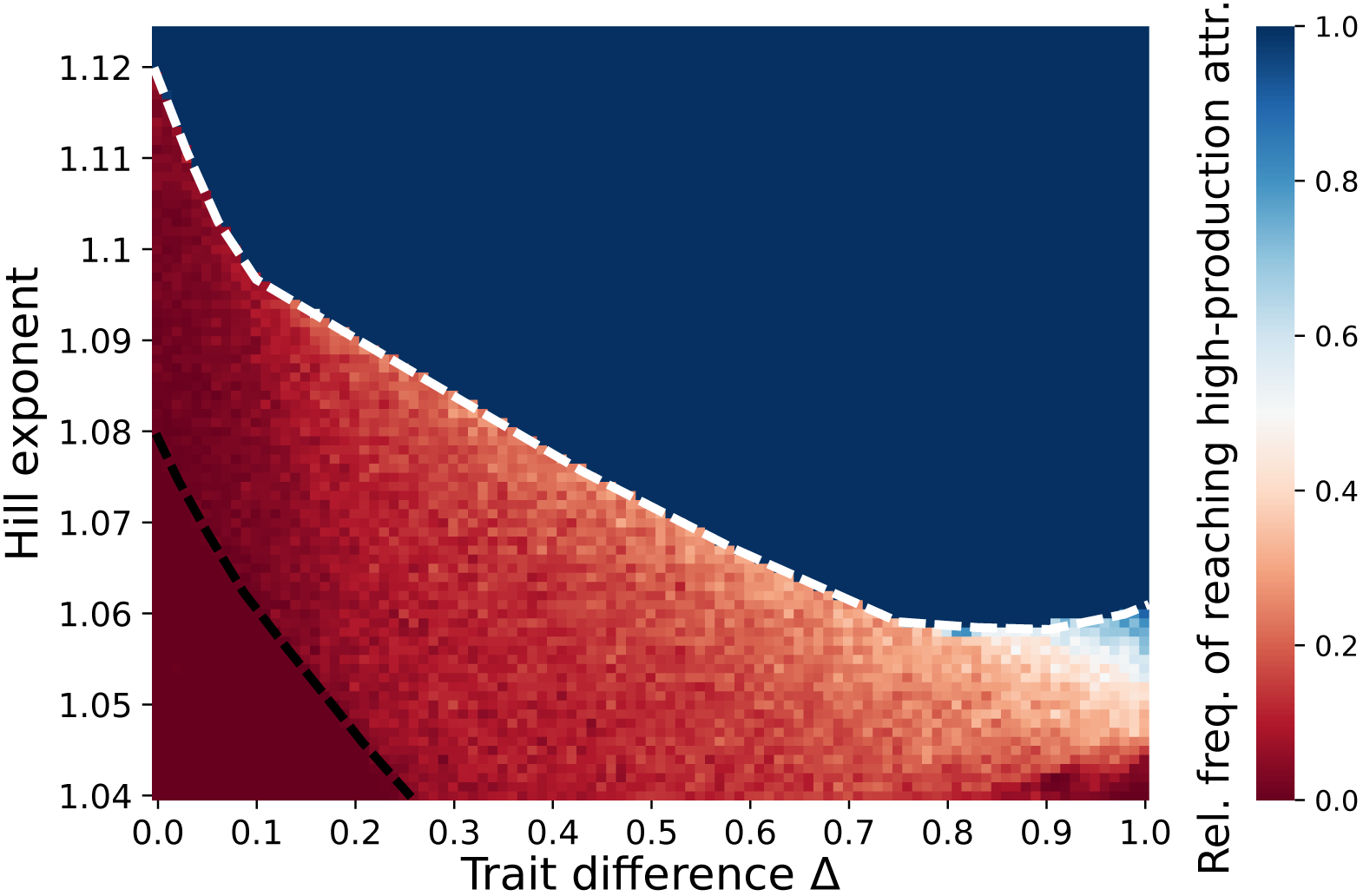
Relative frequency of reaching the high-production state, as a function of the trait difference ∆ and the Hill-exponents h = η. Each of the points in the 101 × 85 grid shows the relative frequency of reaching the high-production state, sampling 200 random initial conditions. The black dashed line shows the approximate location of the boundary crisis of the high-production state. The low-production attractor also undergoes a boundary crisis, the approximate location of which is indicated by the white dashed line.

As trait variation (∆) increases, generalist and specialist consumers at the top level exploit different temporal niches and force the intermediate level to split up into two distinct groups comprising the defended and the undefended species, respectively. As both groups include both generalist and specialist consumers, this further weakens the interaction between the intermediate and basal levels, strengthening the aforementioned mechanisms that stabilize the high-production state. Notably, the mean population biomasses stay relatively constant as trait variation increases (cf. Fig. 6c & 6d), while the community temporal variability decreases (Fig. 6e & 6f, gray lines). This effect can also be predicted from the phase relationship diagrams, which show that with increasing trait variation, competing species within the same trophic level move out-of-phase with each other (Fig. 4c, 4e & 4f). Such out-of-phase cycles indicate compensatory dynamical patterns with potentially high amplitudes at the population level. However, because the different species are able to exploit different temporal niches, the community temporal variability is kept low.

**Figure 6:**
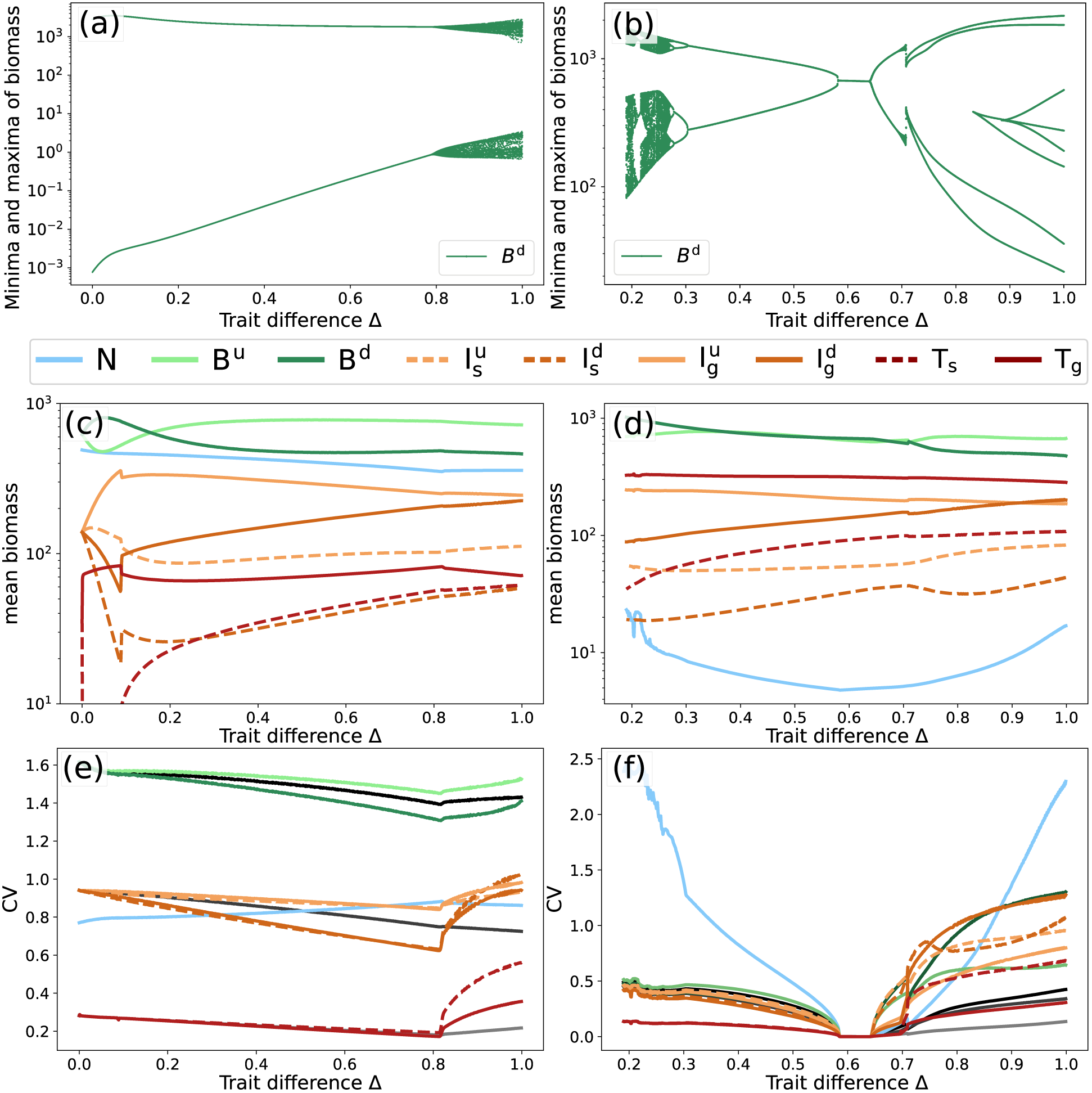
Bifurcation diagrams of the defended basal species, B^d^ ((a), (b)), mean biomasses ((c), (d)), and CV ((e), (f)) of all the species in the system, for the low-production (left) and the high-production (right) attractors, for h = 1.05. For the bifurcation diagrams of the other species see Appendix C. The CV of the specialist top predator is not plotted for the region where it goes extinct (0 ⪅ ∆ ⪅ 0.1 in panel (e)). In panels (e) and (f), the black to gray lines respectively denote the CV of the first, second, and third trophic level as a whole. Note the different scales on the x-and y-axes.

In summary, adding trait variation safeguards the high-production state which is characterized by a high top-level biomass resulting from an efficient transfer of energy towards the higher trophic levels, and low temporal variability due to weak coupling between the intermediate and basal levels and prominent compensatory dynamics within the lower trophic levels. Contrarily, losing trait variation increases the risk of an irreversible transition to the low-production state, which is characterized by a lower top-level biomass resulting from the less efficient transfer of energy towards the higher trophic levels, and higher temporal variability. With trait variation added, primary production increases from the low-to the high production state by a factor of 1.5 and the efficiency of the energy transfer towards the top level increases by a factor of 2 (See Table C1 in Appendix C). Hence, adding trait variation results in a more productive and energy-efficient food web.

More details about the above-mentioned results are presented in the respective sections below.

### 3.1 Phase relationships as a way to identify underlying mechanisms

In order to understand how to use the phase relationships between different populations of a complex food web such as the maximally trait-separated web (Fig. 1c) to uncover the mechanisms driving their dynamics, let us first look at the simpler linear chain containing only three species (Fig. 1a). As mentioned above, the state shown in Fig. 3a with a mean nutrient concentration of about 250 *μg* N/*l* will be called the low-production state, relative to the other state (Fig. 3b) which has a much lower mean nutrient concentration of around 10 *μg* N/*l* and therefore will be called the high-production state.

Closer inspection of the time series reveals the origin of the difference in mean free nutrient levels between the two states. In the low-production state (Fig. 3a) the intermediate level is able to grow to sufficiently high densities to graze the bottom level down significantly, despite the predation pressure imposed by the top species. Hence, the nutrient uptake is strongly reduced for a considerable amount of time leading to a relatively high mean nutrient level. Conversely, in the high-production state (Fig. 3b), the higher biomass at the top level implies a stronger control over the intermediate level. The intermediate species are thus not able to grow to the density levels reached on the low-production state, and in turn, do not graze the basal level down to low densities. Hence, the mean nutrient level is much lower. From this, we could conclude that the overall control exerted by the top level is higher in the high-production state than on the low-production state. Such an observation cannot be made as straightforwardly by inspecting only the mean biomass levels, as the temporal averages of the intermediate and basal biomasses are quite similar in both states and thus, they do not inform about potential changes in the production at each level. Therefore, examining the degree of top-down or bottom-up control in the case of non-static dynamics requires information about the oscillations themselves.

Interestingly, the phase differences between the different trophic levels contain sufficient information to reach the same conclusions regarding the strength of top-down control in the two states. In the low-production state, the phase differences between the top and intermediate level, and intermediate and bottom level, are about a ¼-cycle Fig. 3c), reflecting the presence of clear predator-prey oscillations, i.e., cyclic change between top-down and bottom-up control, between both the top and intermediate level, and the intermediate and basal level. In contrast, in the high-production state, the phase lag between the intermediate and basal level is significantly more than a ¼-cycle (Fig. 3d), indicating rather weak interactions between these two different trophic levels.

With this in mind, we now investigate the fully trait-separated food web (∆ = 1, Fig. 1c), whose dynamics are shown in Fig. 4a and 4b. Just as the linear chain, the system settles down to a stable limit cycle. While the dynamics are visually much more complex when compared to the linear chain, the basic properties and differences between the two states remain the same. However, in contrast to the chain, the discrete Fourier frequency spectra (cf. Appendix C, Fig. C3) reveal two distinct frequencies at substantially different timescales. Despite this increase in complexity, our results clearly show that the phase relationships between distinct populations of adjacent trophic levels provide substantial information about the regulations of trophic interactions and changes therein.

The low-production state of the maximally separated food web (Fig. 4a) exhibits two important timescales governing the overall dynamics. First, the same high-frequency oscillations as were observed for the food chain are present, with the ¼-lag cycles indicative of predator-prey oscillations (Fig. 4d). Second, oscillations on a slower timescale are found. Their phase relationships show that they arise from the trait differences between species (Fig. 4c). Here, the top species are almost completely out of phase relative to each other. Consider first the specialist predator *T*_s_, which preys only on the undefended intermediate species 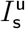 and 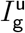. The phase relationship diagram shows that these species precede *T*_s_ by the regular 1/4-lag. The same is true for *T*_g_, which is preceded by a quarter-cycle by the defended intermediate species 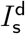 and 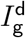. Quarter-lag cycles are not observed between the basal and intermediate trophic level, which indicates that the trait differences within the top trophic level influence the intermediate level more strongly as those on the intermediate level influence the basal level. As the two alternating groups of defended and undefended intermediate species contain a general and specialist grazer on the basal species, they exert together approximately the same grazing pressure on both types of basal species. Consequently, no clear phase relationship between the intermediate and basal level is found. However, visual inspection combined with analysis of the Discrete Fourier Transform (DFT) spectrum (Appendix C, Fig. C3) shows that the high-frequency component is the dominant one, explaining most of the observed variation in the biomass. Hence, the biomass dynamics reflect an overall balance between top-down and bottom-up interactions in the low-production state, similar to the simple linear chain.

In the high-production state of the maximally separated food web (Fig. 4b), the difference in dynamics as compared to the linear chain is even more pronounced. The basal species exhibit a clear compensatory dynamical pattern, with alternating biomass peaks of defended and undefended species. While the dynamics appear highly irregular, the frequency spectrum shows that they are also mainly driven by two frequencies. On the lower of these frequencies, which explains most of the variation observed in this state, the phase relationships resemble those of the low frequency in the low-production state (Fig. 4e vs. 4c), with the exception of the basal species, which now cycle out of phase. The specialist and generalist top species also move out of phase, which leads the groups of defended and undefended intermediate species to behave similarly, as they each precede their respective main predator. As in the low-production state, no further relationship can be identified between the intermediate and the basal level.

However, the high frequency roughly corresponding to that of the chain also has an influential component in the Fourier spectrum. On this frequency, the phase relationships show that the basal species also move out-of-phase. In contrast to the dominant lower frequency, the intermediate species are now split into two groups according to their main prey type. The generalist intermediate species follow the defended basal species, and the specialist intermediate species the undefended basal species, by a ¼-lag. As each of these two groups of intermediate species contains both a defended and undefended type, no further relationship can be drawn between the phases of the intermediate and the top level.

In summary, the strength of top-down control across trophic levels can be inferred from the phase relationships in both the linear chain and the maximally separated food web. The phase-relationships further reveal compensatory dynamics within trophic levels in the fully separated web.

### 3.2 Trait variation increases resilience of the high-production state

Recall that trait differences between the modelled species at each trophic level, determined by ∆ (Eq. (8)), can be varied continuously. Varying ∆ between ∆ = 0 (chain) and ∆ = 1 (maximally separated web) reveals the intermediate region between the two extremes considered so far.

In this intermediate region, the food web is not yet completely separated as is the case for Fig. 1c, although there are already trait differences between different species at each trophic level. That is, the specialist predator species are not yet fully specialized: they are still able to prey on the defended species albeit with a lower efficiency than the undefended species. Accordingly, the undefended species are not fully defended against the specialist predators. The difference in growth rates between defended and undefended species is thus gradually increased to its maximum value, which is obtained when ∆ = 1. In this way the trade-offs between defense and growth rate, and between specialization and prey grazing efficiency, are explicitly built into the model.

To investigate the effect of trait variation on the likelihood of the system adopting either of the two alternative stable states, we determined the size of the basin of attraction of the high-production state. Fig. 5 shows the relative frequency of a random initial value falling in this basin of attraction, as ∆ is varied. The random initial values were sampled from the set of all potentially accessible biomass configurations of the chemostat system, i.e., the total carbon content in the system does not exceed the maximum possible carbon content attainable by the incoming nutrient concentration *N*_0_.

The region of intermediate frequency values confirms that the bistability is an important aspect of the system. The typical behavior when varying ∆ from 0 to 1 is an increasing probability of reaching the high-production state. Furthermore, the graph shows an important dependency on the predator-prey functional responses’ Hill coefficient *h*: increasing the exponent also increases the probability of reaching the high-production state. Investigating the effect of other model parameters on the presence of bistability reveals that it is quite common for this type of model structure, and that the trends presented here are not limited to this particular part of the parameter space. For details, see Appendix B.

Over the whole range of ∆, there is a very sharp transition between the region where only the high-production state exists (dark blue), and the region where both states exist, indicated by the white dashed line (Fig. 5). Notably, the border decreases steeply as the trait difference ∆ is increased, indicating the much lower dependence on low-density grazing suppression for higher amounts of standing trait variation. The sharp border between the two regions is an indication that the low-production state undergoes a catastrophic bifurcation, where it suddenly disappears. Similar behavior is observed for the high-production state, indicated by the black dashed line. This transition is of particular ecological interest as it implicates the sudden disappearance of the high-production, low-temporally-variable state. The region for *h <* 1.04 was not considered, as the amount of extinctions was too high. However, the graph indicates that the probability of reaching the high-production state decreases further.

### 3.3 Dynamical properties of the alternative stable states under gradual changes in trait variation

Consider now *h* = 1.05 as a representative value catching the most complex region in Fig. 5, to study the possible effects of varying ∆ on the system’s dynamics. In this case, the low-production state exists over the whole range of ∆, its the bifurcation diagram for the defended basal species *B*^d^ is shown in Fig. 6a. The qualitative features of the diagram are representative for the other species in the network, whose bifurcation diagrams are shown in Appendix C, Fig. C1 & C2.

For ∆ = 0, the oscillations are simple, in the sense that they are governed by a single timescale and have a constant amplitude, as the maxima and respectively minima each fall on the same position. As ∆ increases, this situation remains unchanged, up until the species become different enough for the second timescale to emerge, in which they exhibit a compensatory dynamical pattern. This explains the variation of the values of the extrema as the two timescales interact destructively and constructively.

The mean biomasses of the species (Fig. 6c) reveal that the specialist top predator goes extinct for low values of ∆. In this case, the species are too similar to stably coexist due to a lack of niche differences, and the generalist predator out-competes the specialist. However, it quickly recovers as ∆ is increased and the species become more different. While this event causes some disturbances in the mean biomasses of the other species, outside of this range the values are more or less constant.

Disregarding the initial region of ∆ where the specialist top predator goes extinct, all species’ *CV* exhibit a gradual decrease as ∆ is increased, up to the point where the second timescale enters the system (Fig. 6e). At this point, a sharp increase is observed as the complexity is enhanced by the interaction of the two timescales. The black to gray lines, depicting the *CV* of the biomasses at each trophic level as a whole, show that the sharp increase is not present on the trophic-level-scale. Hence, the increase in *CV* for each of the species can be solely attributed to the introduction of the second, slower timescale. As discussed above, species with different traits may move out of phase on this timescale, and thus the effects of the slower timescale on the temporal variability for the trophic level as a whole cancel out.

The bifurcation diagram for the high-production attractor (Fig. 6b) does not cover the full range of 0 *≤* ∆ *≤* 1 for *h* = 105, as it only exists on the right side of the black dashed line in Fig. 5. Furthermore, the attractor exhibits a much richer structure as ∆ is varied than the low-production attractor. Multiple bifurcations occur in which the dynamics are altered. In particular, lowering ∆ sufficiently the system undergoes a series of period-doubling bifurcations which lead to chaotic dynamics. Eventually the attractor undergoes a boundary crisis, as indicated by both the sudden disappearance of the then chaotic attractor and the presence of a chaotic transient (Appendix C, Fig. C4).

The species’ mean biomass on the high-production state (Fig. 6d) reveal a similar monotonicity as those on the low-production state (Fig. 6c). A notable observation is the very low mean nutrient level along the whole range of ∆. The nutrients show a very high *CV* (Fig. 6f), which can be attributed to their low mean value. In addition, the *CV* for each of the species is higher on the right side of the Hopf-bifurcations (higher ∆), as compared to the left side for lower values of ∆. However, just as for the low-production state, these increases are buffered when looking at the temporal variability of the trophic levels as a whole (black to gray lines). This reflects the compensatory dynamical pattern of the high-production state, where some of the species move out of phase, which leads to a reduction in temporal variability on the entire trophic level.

## 4 Discussion

We developed a generic tritrophic model to investigate the effect of varying degrees of trait variation on the dynamics of multitrophic food webs and their associated ecosystem functions such as the mean resource use efficiency, biomass production, temporal variability and resilience. By increasing the trait difference parameter ∆ from 0, the system increases in complexity while it changes gradually from a simple chain without trait variation—where all species are generalists—to a complex web with generalist and specialist consumers, and correspondingly defended and undefended prey. The relevant parameters affecting these traits (growth rate, edibility, food preference, and half saturation constant) are closely linked to the functions of the individual species in the food web. Hence, increasing ∆ also increases the functional differences between the species, and thus, the functional diversity of the system. For ∆ *>* 0 but low, the trait differences are small which means the species are very similar, hence, the functional diversity at each trophic level is low. Correspondingly, for ∆ close to one, the functional diversity of the system is high, even though the number of species is kept constant. Therefore, varying ∆ is a means to study the effects of changing functional diversity on all three trophic levels on the dynamics of the whole system without potentially confounding effects of changing the number of species. The different aspects of how trait variation impacts the food web dynamics are discussed in detail below.

### Phase relationships help unravel complex trophic interactions

Traditionally, effects of multi-trophic interactions such as trophic cascades and the degree of bottom-up or top-down control were studied using a rigid linear chain in equilibrium (Carpenter et al., 1985; Pace et al., 1999). However, natural systems are usually not simple chains, but highly complex webs with functionally diverse species at all trophic levels (Boit et al., 2012; Wollrab et al., 2012). Moreover, their dynamics may not evolve towards an equilibrium fixed point, but rather to a limit cycle (May, 1972), or a strange attractor (Hastings et al., 1993) where they will perpetually exhibit oscillatory behavior. This phenomenon can be separated from stochastic noise and has been observed in natural communities (Kendall, 1998). Such oscillatory behavior gives rise to certain phase relationships between the biomass dynamics of the different species.

Additionally, in the maximally trait-separated food web (Fig. 1c), calculation of the Discrete Fourier Spectrum clearly exposes the two timescales at which major driving mechanisms take place. The emergence of a second timescale does not rely on the addition of a third trophic level as this feature has already been found in bitrophic models that considered multiple species or phenotypes at only one (Yamamichi et al., 2011) or both trophic levels (Tirok and Gaedke, 2010). However, our treatment highlights how the phase relationships may shed light on the mechanisms driving complex systems by disentangling the different timescales at which these mechanisms may act.

#### Strength of trophic interactions

We found that the main dynamical differences between the two alternative stable states present in our system can be explained by an increased top-down control of the top level on the intermediate level. When the intermediate level is strongly controlled, such as is the case on the high-production state, its species are unable to control the basal level. The basal level is in turn able to fully exploit the available nutrients, increasing the overall production in the system (See Appendix C, Table C1).

This result holds independent of the amount of trait variation present, and is in line with previous studies showing that reduced top-down control may result in an increased phase difference between predator and prey (Yoshida et al., 2003; Becks et al., 2010). Importantly, the larger than ¼-cycle phase difference between the basal prey and intermediate predator observed in our system with only one species per trophic level (Fig. 1a) shows that the common conception of anti-phase cycles as a “smoking gun” for the presence of evolution, or other mechanisms causing trait changes (Ellner and Becks, 2011; Hiltunen et al., 2014) does not hold any longer when considering multitrophic systems in which the intermediate predator faces strong top-down control by the top predator.

#### Role of compensatory dynamics

When a community consists of functionally diverse populations, a decline in one functional group can be accompanied by an increase of another (Klug et al., 2000). In this way, even though the individual populations exhibit high temporal variability in their biomasses in our model, the variability of the community biomass per trophic level remains low (Fig. 6f). Such an effect has been observed before in studies investigating the effect of standing trait variation or phenotypic plasticity on population dynamics (Kovach-Orr and Fussmann, 2013; Bauer et al., 2014), and it is often made possible through compensatory dynamics between the species (Micheli et al., 1999; Vasseur and Gaedke, 2007; Gonzalez and Loreau, 2009). Hence, compensatory dynamics can be understood as a mechanism by which ecosystem functions such as biomass production can stay rather constant while individual populations may be highly variable (Hooper et al., 2005; Bauer et al., 2014). Compensatory dynamics are observed in both the high-and low-production state, for sufficiently high ∆ (Fig. 4a & 4b). When present, they effectively decrease the biomass *CV* of the trophic level as whole, even though the *CV* s of the species’ individual biomass may be relatively high (Fig. 6e & 6f). These compensatory dynamical patterns naturally keep species within a trophic level moving out-of-phase relative to each other, and thus, can also be inferred by analyzing phase relationship diagrams.

Notably, the compensatory dynamics on the low-production state are only present at the slower timescale related to the trait dynamics (Fig. 4c). For example, the dominant faster timescale does not exhibit compensatory dynamics (Fig. 4d), and thus, given substantial variation in the biomass of the individual populations, the *CV* at the community level remains relatively high (Fig. 6e). Even so, the sharp increase in temporal variation on the population level for high trait variation is buffered on the community scale, through the compensatory dynamics taking place on a different timescale than the dominant one. Our time-scale dependent phase-relationships between populations are in line with empirical observations showing that phytoplankton populations may exhibit compensatory dynamics on the sub-annual scale, likely associated with trophic interactions, combined with synchronous dynamics on the annual, externally driven timescale (Vasseur and Gaedke, 2007). Similarly, zooplankton dynamics may be governed by two distinct timescales: seasonal variation and experimentally varied environmental conditions (Keitt and Fischer, 2006). Hence, unraveling the different timescales governing the population dynamics may help to understand the major processes driving them.

### Trait variation promotes high production at the top level

In line with our results, bistability has been observed in other food chain models (Abrams and Roth, 1994; Letellier and Aziz-Alaoui, 2002; Van Voorn et al., 2010; Erbach et al., 2013), ontogenetic growth models (Guill, 2009; Nakazawa, 2011), and in other, broader ecological contexts (Beisner et al., 2003). The presence of two alternative states in our system is an important feature as it may have far-reaching consequences regarding the stability and perseverance of food webs when confronted with external perturbations. A commonly made distinction when studying the effects of perturbations is whether they consist of a change to the state variables, or to the actual model parameters (Bender et al., 1984; Beisner et al., 2003). The first kind, for example a sudden decrease in one of the species’ biomass, is often called a pulse perturbation because of its short duration. The second kind is called a press perturbation, because the change to the perturbed parameters is permanent, such as a decrease in the nutrient inflow concentration. In a multistable system, pulse perturbations, particularly when they are large, might push the system over the edge of one basin of attraction into another, where the dynamics are potentially completely different. Press perturbations may produce a similar outcome by causing large changes to an attractor’s basin of attraction, or by crossing a bifurcation point where the dynamics change significantly. Therefore, the size of the basin of attraction may be used as a measure of resilience (Beisner et al., 2003). A highly resilient system will nearly always return to its original state, hence its basin of attraction must be very large. Conversely, a non-resilient or fragile system is easily pushed out of one basin of attraction into another one.

Recall that the two states in our system have very different dynamical properties: the low-production state with low top biomass production and high variability, compared to the high-production state with high top biomass production and low variability. From an ecosystem function perspective, low variability or high biomass in higher trophic levels are beneficial for e.g. fish yield. Therefore, it may be desirable to keep the system on the high-production state.

While biomass production of a community is known to be mostly positively correlated with its functional diversity (Tilman et al., 1997; Naeem et al., 2012), we also found the high-production state in the food chain. This corresponds to, e.g., modern agricultural systems, which typically consist of monocultures with a low functional diversity, but a high biomass production available for higher trophic levels. However, even though such monocultures may produce more biomass than some functionally highly diverse mixtures, they are very fragile against external disturbances (Yachi and Loreau, 1999; Loreau et al., 2001). In this way, functional diversity is regarded as an insurance against external perturbations. We clearly observed such an effect in our system, for both pulse and press perturbations, as illustrated by Fig. 5. Since the basin of attraction of the high-production attractor increases in size with ∆, the system becomes less likely to be pushed out of the basin of attraction by a pulse perturbation. This trend is persistent when varying not only the Hill exponents, but also the dilution rate, and the nutrient inflow concentration (Appendix B, Fig. B1b & B1a), and is thus not limited to a particular part of the model’s parameter space. In addition, the boundary crisis causing the sudden disappearance of high-production attractor (Fig. 5, black dashed line) is only present for low values of ∆. Hence, functional diversity also protects the high-production state from suddenly disappearing under a press perturbation.

Typical for a boundary crisis, as the high-production state undergoes when decreasing ∆, are the long transients that are still present near the crisis point (Grebogi et al., 1982) (Appendix C, Fig. C4). In an ecological context this could be problematic as such a long transient implies there is no way to know exactly when the crisis point has been passed and the basin of attraction no longer exists, until the system eventually accelerates towards the only remaining attractor. Such regime shifts were empirically observed and predicted by a variety of ecosystem models in different contexts (Scheffer and Carpenter, 2003), such as woodlands threatened by fires turning into grasslands (Dublin et al., 1990), and shallow lakes threatened by eutrophication turning from a macrophyte to a phytoplankton dominated state (Scheffer et al., 1993). The key idea is that a small perturbation near the bifurcation point may move the system to an alternative stable state, but once this has happened, a much larger perturbation is needed in order to return back to the original state. It has been argued that, under certain circumstances, one may be able to observe early-warning signals that a transition is imminent (Scheffer et al., 2009; Carpenter et al., 2011; Kéfi et al., 2014). For example, near some types of bifurcations a dampening of the speed-of-return after a pulse perturbations may be observed, called critical slowing down (Wissel, 1984; Scheffer et al., 2009). In the case of a boundary crisis, showing the existence of any early-warning signals has proven to be difficult (Hastings and Wysham, 2010; Boettiger and Hastings, 2012). However, even if their existence could be shown mathematically, they will almost certainly be very difficult or impossible to detect in a real-life setting, where the exact chaotic dynamics may be obscured by measurement noise. Our results reveal that maintaining sufficient trait variation provides protection from boundary crises, with their often ecologically and economically undesirable consequences.

These conclusions rely on the presence of alternative stable states in our model. This is a prominent property in tritrophic systems, present in even a simple tritrophic chain with Holling-type-II functional responses and logistic growth of the basal species (Abrams and Roth, 1994). However, there always exist parameter regions where there is only one non-trivial stable state. We find that also in such cases, production at the top level and temporal stability both increase with ∆, as the attractor changes from resembling the low-production to resembling the high-production state in a gradual way (See Appendix B, Fig. B4).

### Influence of a sigmoidal functional response

The use of sigmoidal functional responses such as the (generalised) Holling type-III (*h* = *η >* 1) has been an active area of discussion for quite some time. Sigmoidal functional responses are praised for their favorable effects on food web dynamics such as an increased dynamical stability (Williams and Martinez, 2004; Kalinkat et al., 2011). Such an increase is justified by the apparent discrepancy between the observed stability of natural ecosystems, and the highly unstable nature of ecosystem models describing them (McCann, 2000). While experimental evidence has traditionally mainly supported hyperbolic functional response shapes, such as Holling type-II (*h* = *η* = 1) (DeMott, 1982; Murdoch et al., 1998), sigmoidal functional responses such as Holling-type-III provide models with additional stability which may overcome this discrepancy. Recent experiments studies have found evidence for sigmoidal functional response shapes (Sarnelle and Wilson, 2008; Morozov, 2010; Kalinkat et al., 2013), or otherwise have shown the difficulty in distinguishing Holling type-II from type-III functional response shapes (Seifert et al., 2014). Furthermore, sigmoidal shapes account for natural processes not captured by the model such as spatial heterogeneity, refuges, formation of resting shapes, etc. Hence, Hill exponents close to—but higher than—one are likely to be relevant, and thus, the requirement of at least some grazing suppression at low densities for all species to coexist adds to the realism of the model. Even in the highly-controlled environment of the chemostat, some of the proposed mechanisms giving rise to the predation dampening at low prey densities, such as prey clumping (Oaten and Murdoch, 1975) or other induced defenses (Lurling and Beekman, 2006), may well be of importance. In addition, while a Hill exponent larger than 1 does facilitate coexistence, it is not a guaranteed. For example, Fig. 6c shows that one of the top predators is not able to survive for 0 ⪅ ∆ ⪅ 0.1.

Nonetheless, Fig. 5 also shows a significant, decreasing dependence on low prey density grazing suppression in order to reach the high-production state. For most of the range of ∆, the sharp border between the bistable region and the region where only the high-production state exists occurs at a lower value of the Hill-exponent as ∆ is increased. Hence, while the grazing suppression at low prey densities is necessary to reach the high-production state, it becomes a less important factor as ∆ is increased.

### Concluding remarks

Despite the higher dynamical complexity of the resulting food web, the introduction of trait variation at all trophic levels to a linear food chain increased the overall reliability of ecosystem functions, such as resource use efficiency and high biomass production. Our results highlight that functional diversity on different trophic levels can reduce the overall temporal variability at the community level through compensatory dynamics among functionally different species within a trophic level. Investigating the phase relationships between the different species of adjacent trophic levels enabled us to identify the regulation of trophic interactions, such as changes in top-down or bottom-up control, in oscillatory dynamical regimes. Accordingly, we observed that strong deviations from the expected ¼-lag between predator and prey are possible in a tritrophic system, even without any trait variation. Hence, observation of such deviations do not necessarily indicate the presence of eco-evolutionary dynamics as is often assumed. Furthermore, independent of the presence or absence of trait variation, our tritrophic model shows two alternative states with the top predator exhibiting either a relatively low or high biomass. However, while the high-production state is attainable in a tritrophic food chain, its basin of attraction is very small. It becomes more resilient when trait variation is added, underlining the role of functional diversity as an insurance against sudden pulse perturbations. In addition, as trait variation decreases, this state may suddenly disappear through a boundary crisis. Hence, high functional diversity also protects the high-production state under press perturbations. We thus highlight the importance of functional diversity regarding resilience against external perturbations, low community temporal variability, resource use efficiency, and maintenance of biomass in higher trophic levels.

## Acknowledgments

We thank A. Boit, E. van Velzen, M. Sieber, M. Raatz and E. Ehrlich for helpful comments and suggestions during the project. This project was funded by the German Research Foundation (DFG) Priority Programme 1704: DynaTrait (GA 401/26-2).

